# Phosphoinositide-focused CRISPR screen identifies novel genetic vulnerabilities in PDAC and AML cells

**DOI:** 10.1101/2024.09.09.612071

**Authors:** Daniel KC Lee, Ryan Loke, Jonathan TS Chow, Martino M Gabra, Leonardo Salmena

**Affiliations:** Department of Pharmacology & Toxicology, University of Toronto, Toronto, Ontario, Canada

**Keywords:** Phosphoinositides, CRISPR-Cas9, AML, PDAC, Functional Screen

## Abstract

Phosphoinositides (PIs) are minor but essential phospholipids that play crucial roles in cellular signaling pathways, membrane dynamics, and the regulation of various cellular processes. We developed and utilized a novel PI-focused CRISPR gRNA library to perform negative-selection and positive-selection screens in PANC-1 and OCI-AML2 cells, models of pancreatic ductal adenocarcinoma (PDAC) and acute myeloid leukemia (AML), respectively. Through these screens, we identified 28 essential genes in PANC-1, 84 essential genes in OCI-AML2, and 28 regulators of colony formation in OCI-AML2. Our results using this small and focused library uncovered false negatives and subtle effects that may be missed in genome-wide approaches, while enabling adaptation to different screening conditions. Overall, our results uncovered previously uncharacterized essential genes in PDAC and AML that can be leveraged as therapeutic targets and biomarkers. We also demonstrate that focused libraries offer a more efficient and targeted approach to uncovering critical genetic determinants of cancer progression.

## Introduction

Phosphoinositides (PIs) are membrane-bound lipid second messengers that play essential roles in fundamental cellular processes, including cell growth, survival, migration, and differentiation.^1^ Signaling in these pathways is tightly controlled by the reversible phosphorylation at positions 3, 4, and 5 of the inositol ring by kinases and phosphatases.^1^ This leads to the creation of different PI species that act as docking sites for a variety of effector proteins to regulate downstream signaling cascades.^1^ Importantly, dysregulation of PI signaling can contribute to the development and progression of cancer, especially through the PI-3-kinase (PI3K)/Akt pathway, which is hyperactivated in >50% of pancreatic ductal adenocarcinoma (PDAC) and acute myeloid leukemia (AML) patients.^2–7^ However, researchers have yet to conduct a comprehensive investigation of the entire PI signaling system beyond examining kinases and phosphatases.^8,9^

Recent advances in CRISPR-Cas9 technology have enabled genome-wide screens to identify essential genes and regulatory pathways in cancer cells.^10^ However, these broad approaches may miss subtle yet significant genetic vulnerabilities and may not be feasible under diverse screening conditions.^10^ Focused libraries offer a promising alternative, providing increased coverage and resolution for curated gene sets.^10^

In this study, we developed and utilized a novel PI-focused CRISPR gRNA library to perform both negative-selection and positive-selection screens in PDAC and AML cell models, specifically PANC-1 and OCI-AML2 cells. Our screens aimed to comprehensively investigate the PI signaling system, encompassing kinases, phosphatases, and effector proteins, to uncover novel regulators of cell growth and colony formation. By identifying essential genes and regulators specific to these cancers, our approach revealed potential therapeutic targets and biomarkers, offering new insights into the molecular mechanisms driving PDAC and AML.

## Materials and Methods

### Cell Line Maintenance

OCI-AML2 cells were maintained in AMEM (Wisent Inc., 310-010-CL) supplemented with 10% FBS (Wisent Inc., 080150) and 1% penicillin/streptomycin (Wisent Inc., 450-201-EL). HEK293T and PANC-1 cells were maintained in DMEM (Wisent Inc., 319-005-CL) supplemented with 10% FBS and 1% penicillin/streptomycin. Cells were dissociated using 0.25% Trypsin-EDTA (Wisent Inc., 325-043-EL). All cell lines were maintained at 37°C and 5% CO_2_. Cell lines were verified to be mycoplasma-free using the PlasmoTest Mycoplasma Detection Kit (InvivoGen, rep-pt1).

### Lentivirus Production

9 million HEK293T cells were seeded in 15 cm dishes and transfected 24 hours later with 50 μg lentiviral vector, 37.5 μg packaging vector psPAX2 (Addgene Plasmid #12260), and 15 μg envelope vector pMD2.G (Addgene Plasmid #12259) using the calcium phosphate method. 24 hours post-transfection the media was replaced, and virus-containing media was collected 48- and 72-hours post-transfection. Cells and debris were removed by centrifugation at 500 x g for 10 minutes and filtration through a 0.45 μm filter. Virus was concentrated by ultracentrifugation, resuspended in PBS (Wisent Inc., 311-010-CL), aliquoted, and stored at −80°C for future use.

### Cas9 Cell Line Generation

3x10^5^ OCI-AML2 or PANC-1 cells were infected with either lentiCas9-Blast (Addgene Plasmid #52962) or pCW-Cas9-Blast (Addgene Plasmid #83481) virus in the presence of 8 μg/mL protamine sulfate (MP Biomedicals, 02194729-CF) for 48 hours in a 6-well plate. After a 24-hour recovery period, infected cells were selected with 10 μg/mL blasticidin (BioShop, BLA477) for 7-10 days. Once stable cell lines were generated, clones were isolated by limiting dilution for PDAC cell lines and colony formation for AML cell lines in a 96-well plate. Clonally derived Cas9 cells were validated by western blot and an EGFP reporter assay to assess Cas9 expression and activity, respectively.

### Assessment of CRISPR-Cas9 Editing Efficiency

3x10^5^ parental and clonally derived Cas9-expressing OCI-AML2 or PANC-1 cells were infected with pXPR_011 (Addgene Plasmid #59702) virus as described above. After a 24-hour recovery period, infected cells were selected with 2 μg/mL puromycin (BioShop, PUR333) for 48 hours and analyzed by flow cytometry either five days post-selection for Cas9 stable lines or 5-days post-induction with 1 μg/mL doxycycline (Thermo Scientific Chemicals, 10592-13-9) for Cas9 inducible lines. The pXPR_011 plasmid contains both EGFP and a gRNA for EGFP. Parental cells that do not express Cas9 are EGFP-positive, while Cas9-expressing cells are EGFP-negative.

### PI-focused CRISPR gRNA Library Design

PI signaling genes (kinases, phosphatases, effectors) were selected from canonical signaling pathways and an extensive literature review. The top 100 core essential and nonessential genes were selected based on BF rank order and CCLE expression data in AML and PDAC cell lines.^11,12^ Possible gRNAs were designed through the Broad Institute GPP sgRNA Designer (https://portals.broadinstitute.org/gpp/public/analysis-tools/sgrna-design). gRNAs were selected for the library in a six-step process. First, we excluded gRNAs that contained poly-T sequences as these sequences act as a termination signal for RNA polymerase III.^13^ Next, a gRNA GC content of 40-80% was accepted. Third, gRNAs were restricted to 5-65% of the protein coding region to avoid target sites that code for amino acids near the N’- and C’-terminus. Next, gRNAs that contained, or by cloning would create a BsmBI restriction site were excluded. Fifth, gRNAs with a Rule Set 2 score less than 0.2 were excluded. Lastly, gRNAs with a Match Bin I score greater than 3 were excluded. After filtering, 228,450 candidate gRNA sequences remained and the top 7 gRNAs by Combined Rank were selected for each gene. 125 non-targeting gRNAs were added to the library. There are 10,863 gRNA sequences in the PI-focused CRISPR gRNA library.

### Pooled gRNA Cloning

gRNAs were synthesized as 73mer oligonucleotides on a microarray (CustomArray) at a density of ∼12,000 sequences and amplified by PCR (Invitrogen, 12344040) as a pool (oligonucleotide template and primers are listed below). The PCR products were treated with T4 PNK (NEB, M0201) for 2 hours at 37°C and subsequently purified using the QIAquick PCR Purification Kit (Qiagen, 28106). The modified lentiGuide-puro vector (Addgene Plasmid #52963) (named lentiGuide-puro-P2A-GFP) was digested with BsmBI (NEB, R0580) at 55°C overnight, treated with Antarctic Phosphatase (NEB, M0289) for 1 hour at 37°C, and gel purified using the QIAquick Gel Extraction Kit (Qiagen, 28704). The library PCR pool was cloned into lentiGuide-puro-P2A-GFP by Gibson Assembly (QuantaBio, 95190). In a single reaction, 0.2 pmol of insert was cloned into 0.04 pmol of digested vector (in a 1:5 vector:insert molar ratio) for 1 hour at 50°C. The Gibson product was diluted 1:5 in nuclease-free water and 1 μL was transformed into 25 μL of Endura electrocompetent cells (Endura, 60242). Four parallel transformations were performed using the assembly product, and the outgrowth media was pooled and plated onto four LB-ampicillin (100 μg/mL) plates (245-mm^2^ Bioassay Dish). Colonies were scrapped off the plates, pooled, and the plasmid DNA was extracted using the QIAGEN Plasmid Maxi Kit (Qiagen, 12163).

#### Oligonucleotide Template for Microarray

GGAAAGGACGAAACACCG**<u>nnnnnnnnnnnnnnnnnnnn</u>**GTTTTAGAGCTAGAAATAGC AAGTTAAAATAAGGC

Primer Sequences to Amplify the CRISPR Library:

Fwd: TAACTTGAAAGTATTTCGATTTCTTGGCTTTATATATCTTGT**<u>GGAAAGGACGAAACA</u> <u>CCG</u>**

Rev: ACTTTTTCAAGTTGATAACGGACTA**GCCTTATTTTAACTTGCTATTTCTAGCTCTAA AAC**

#### Lentiviral Titer Determination

3x10^5^ OCI-AML2 or PANC-1 Cas9 cells were infected with dilutions of the lentiviral gRNA library as described above. 48 hours after infection, the percentage of GFP-positive cells was determined by flow cytometry. The percentage of GFP-positive cells was used to determine the lentiviral titer of the lentiviral gRNA library. The lentiviral titer was calculated as follows:

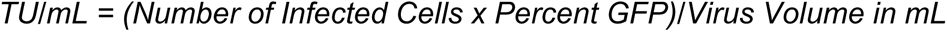

### PANC-1 Pooled gRNA Dropout Screen

40 million clonally selected PANC-1 Cas9 cells were infected with the 11k gRNA library at an MOI of ∼0.3 and 1000-fold coverage as described above. After a 24-hour recovery period, infected cells were selected with 2 μg/mL puromycin for 48 hours. After selection, cells were split into three replicates at 1000-fold coverage and passaged every 3 days at 1000-fold coverage. Cells were collected at 1000-fold coverage for genomic DNA extraction at day 0 (T0) and every 3 days, from day 3 (T03) until day 21 (T21) post-selection. Genomic DNA was extracted from 11 million cells using the QIAamp DNA Blood Midi Kit (Qiagen, 51185), according to the manufacturer’s protocol. Following extraction, the genomic DNA was precipitated using sodium chloride and ethanol and resuspended in buffer EB. NGS library preparation and sequencing were performed by the Princess Margaret Genomics Centre.

### OCI-AML2 Pooled gRNA Dropout Screen

40 million clonally selected OCI-AML2 Cas9 cells were infected with the 11k gRNA library at an MOI of ∼0.3 and 1000-fold coverage as described above. After a 24-hour recovery period, infected cells were selected with 2 μg/mL puromycin for 48 hours and maintained for 7 days post-selection. On day 7, cells were split into three replicates at 1000-fold coverage and passaged every 3 days at 1000-fold coverage. Cells were collected at 1000-fold coverage for genomic DNA extraction at day 7 (T0) and every three days from day 10 (T03) until day 22 (T15) post-selection. Genomic DNA was extracted from 11 million cells using the QIAamp DNA Blood Midi Kit, according to the manufacturer’s protocol. Following extraction, the genomic DNA was precipitated using sodium chloride and ethanol and resuspended in buffer EB. NGS library preparation and sequencing were performed by the Princess Margaret Genomics Centre.

### Analysis of CRISPR Dropout Screen Data

FASTQ files were demultiplexed and trimmed using an in-house Python script. Reads from each sample were mapped to the gRNA sequence library using Bowtie2.^14^ After alignment, read counts for each sequence were tabulated to generate a read counts table. Read counts for each sample were normalized to 10 million reads, and the log_2_ fold change relative to T0 was calculated for each gRNA with at least 30 non-normalized reads in the T0 sample. The Bayesian Analysis of Gene Essentiality (BAGEL) algorithm was used to calculate a BF for each gene using a reference set of essential (n=100) and nonessential (n=100) genes.^11,12,15^ The final BF for each gene is the sum of all gRNA BFs across all replicates at a given timepoint. We chose a BF of 3 and 5% FDR as a cutoff to identify essential genes. The BAGEL software was obtained from https://github.com/hart-lab/bagel.

### Pooled gRNA CFU Screen

20 million clonally selected OCI-AML2 Cas9 cells were infected with the 11k gRNA library at an MOI of ∼0.3 and 500-fold coverage as described above. After a 24-hour recovery period, infected cells were selected with 2 μg/mL puromycin for 48 hours and maintained for 7 days post-selection. On day 7, 1.5x10^5^ cells in 9 mL of null AMEM was combined with 12 mL of 2.1% (w/v) AMEM methylcellulose (ShinEtsu, SM-40C0) and 9 mL of FBS; 10 mL was plated in duplicate on 100x20 mm dishes. Plates were incubated for 7 days at 37°C and 5% CO_2_. Cells that did or did not form colonies were collected by solubilizing the methylcellulose and repeating the procedure two times for enrichment purposes. Cells were collected at 300-fold coverage for genomic DNA extraction at day 7 (T0) and every 7 days, from day 14 (T07) until day 28 (T21) post-selection. Genomic DNA was extracted from 4 million cells using the QIAamp DNA Blood Midi Kit, according to the manufacturer’s protocol. Following extraction, the genomic DNA was precipitated using sodium chloride and ethanol and resuspended in buffer EB. NGS library preparation and sequencing were performed by the Princess Margaret Genomics Centre.

### Analysis of CRISPR CFU Screen Data

FASTQ files were demultiplexed and trimmed using an in-house Python script. Reads from each sample were mapped to the gRNA sequence library using Bowtie2.^14^ After alignment, read counts for each sequence were tabulated to generate a read counts table. Read counts for each sample were normalized to 10 million reads, and the log_2_ fold change relative to T0 was calculated for each gRNA with at least 30 non-normalized reads in the T0 sample. Individual gRNA fold change values were averaged and Z-transformed at the gene-level. Statistical significance was determined by two-tailed, two-sample t-test assuming unequal variance. We chose a |Z-transformed log_2_ (fold change)| > 2, p-value < 0.05, and at least 4 functional gRNAs as a cutoff to identify regulators of colony formation.

### Individual gRNA Cloning

gRNAs targeting *LacZ*, *AAVS1*, *EIF3D*, *ATP6V1H, DCTN2, FUBP1, G6PD, GPN1, IPO7, ITGAV*, *NAA15*, *PACSIN1, S100A9, TPR*, and *UNC45A* were cloned into either lentiGuide-Puro or a modified lentiGuide-Puro plasmid with a GFP cassette.^16^ To clone individual gRNAs, the vector was digested with BsmBI at 55°C overnight, dephosphorylated with Antarctic Phosphatase for 1 hour at 37°C, and gel purified using the QIAquick Gel Extraction Kit. Oligonucleotides coding for the gRNA sequence were phosphorylated using T4 PNK (NEB, M0201) for 30 minutes at 37°C then annealed by heating to 95°C for 5 minutes and cooling to 25°C at 5°C/minute. Phosphorylated/Annealed oligo pairs were ligated into BsmBI digested vector with T4 DNA Ligase (NEB, M0202) and transformed into DH5α cells. DNA was prepped with the QIAGEN Plasmid Maxi Kit and verified by Sanger sequencing. Note that gRNA sequences were chosen from the 11k gRNA library if they were shown to be functional in the screen. All gRNA sequence oligos were designed as follows:

5’ - CACCGNNNNNNNNNNNNNNNNNNNN – 3’
3’ — CNNNNNNNNNNNNNNNNNNNNCAAA - 5’

LacZ: CCCGAATCTCTATCGTGCGG

AAVS1: GATGGTAAGGAGGACTCGAT

EIF3D-1: TGTAGGTTGCCTCCATGGCC

EIF3D-2: AGACGACCCTGTCATCCGCA

ATP6V1H-1: GATGTAGCAAGAACACTGCG

ATP6V1H-2: GGATCCTGGCGATTCAACAT

DCTN2-1: GGAGCTGACAAGCACAAGTG

DCTN2-2: GTAGGGTACCCACTTAGCCA

FUBP1-1: CAAAAATTGGAGGTGATGCA

FUBP1-2: CAAGACGGGCCGCAGAACAC

G6PD-1: AGAGGTGCAGGCCAACAATG

G6PD-2: CTTGCCCCCGACCGTCTACG

GPN1-1: TCCAGGTTGATCACATACGG

GPN1-2: TATGTGTTGATTGACACACC

IPO7-1: ATCGACCTGAGTTACCATGG

IPO7-2: TTAACCAAGTAATCCAGACG

ITGAV-1: AGTTCTCCAATGGTACAATG

ITGAV-2: GCCTTAACAATCAATGTCAG

NAA15-1: GTACTGGACCCAAAGTAATG

NAA15-2: TGTATAGGAAGGAACATCAG

PACSIN1-1: TGGCTTCAAGGAGACGAAGG

PACSIN1-2: TATCACAAGCAGATCATGGG

S100A9-1: CCAGCTTCACAGAGTATTGG

S100A9-2: CTTCACAGAGTATTGGTGGA

TPR-1: AGAGTCTGCGTTATCGACAA

TPR-2: GGAGTGTAAAGCATCTTGGG

UNC45A-1: ACCCTCTGGGTCATCGACCA

UNC45A-2: GGTGTCAAAAAAGGCTTCCG

### Competitive Cell Growth Assay

To validate candidate hits, competition assays were conducted whereby cells expressing gRNA-*LacZ* were co-cultured with cells expressing GFP and a gRNA target sequence, and then were allowed to grow and compete during a 3-week period. In brief, OCI-AML2 tet-Cas9 or PANC-1 Cas9 cells were infected with a virus expressing either lentiGuide-LacZ-Puro or lentiGuide-gRNA-Puro-P2A-GFP as described above. After a 24-hour recovery period, infected cells were selected with 2 μg/mL puromycin for 48 hours, and then gRNA-*LacZ* cells were mixed 1:1 with gRNA-GFP cells in a 6-well plate (75,000 cells per construct). GFP fluorescence was measured on seeding day (T0) and every 3 days for 3 weeks by flow cytometry. In the case of OCI-AML2 tet-Cas9 cells, 1 μg/mL doxycycline was added on seeding day and every 3 days to induce Cas9 expression. Relative GFP fluorescence was calculated by normalizing the percentage of GFP positive cells to T0. Statistical Analysis of three independent experiments (mean±SEM) was assessed using a One-Way ANOVA with Dunnett’s Post-Hoc Test in GraphPad Prism 9.3.0.

### Western Blotting

Cells were lysed in 1X RIPA Buffer (CST, 9806) supplemented with protease inhibitors (Roche, 11836153001) and then centrifuged at max speed for 20 minutes at 4°C. Supernatant was collected and protein concentration was determined using the Bio-Rad Protein Assay (as described on https://www.bio-rad.com/). 20-30 μg of protein was resolved on 10% SDS-PAGE gels in Tris-Glycine running buffer, before being transferred to either nitrocellulose (BioRad, 1620115) or PVDF (BioRad, 1620177) membrane at 60V for 2 hours at 4°C. All membranes were blocked with either 5% BSA or skim milk for 1 hour at room temperature and then washed in TBS with Tween 20. Membranes were blotted with primary antibody overnight at 4°C, washed, blotted with appropriate HRP-conjugated secondary antibodies for 90 minutes at room temperature, and washed again. Blots were exposed using KwikQuant Ultra Digital-ECL Substrate (Kindle Biosciences, R1002) and the accompanying KwikQuant Imager. Antibodies used in this study were: β-Actin (1:10000, CST #4967), Cas9 (1:1000, CST #14697), and GAPDH (1:10000, CST #2118).

### Survival Analysis

Survival analysis with gene expression data from the TCGA-LAML and TCGA-PAAD patient dataset was performed in GraphPad Prism 9.3.0. The gene expression association to survival was plotted using the Kaplan-Meier method and analyzed using the log-rank test. P values < 0.05 were considered statistically significant.

## Results

### Design of the PI-focused CRISPR gRNA Library

Given the lack of comprehensive tools to study the PI signaling system, we aimed to design a PI-focused CRISPR knockout library, named the **PI**ome CRISPR KO library, targeting genes with known or suspected roles in PI metabolism and signaling. To build this library, we first compiled a list of genes involved in PI signaling, including kinases, phosphatases, and effectors, from various studies (**Table S1**).

Among the numerous gRNA design tools at our disposal, we discovered that the Root lab’s tool stands out as the most comprehensive.^17–20^ This tool goes beyond simply predicting gRNA on-target activity; it also provides insightful analyses regarding off-target effects.^19^ By leveraging this tool, we systematically generated all conceivable gRNAs targeting the selection of PI genes. Subsequently, we meticulously sieved through the pool of gRNA candidates using a rigorous six-pass screening methodology (e.g. on-target activity, target location) (**Figure 1A**).

**Figure 1.**
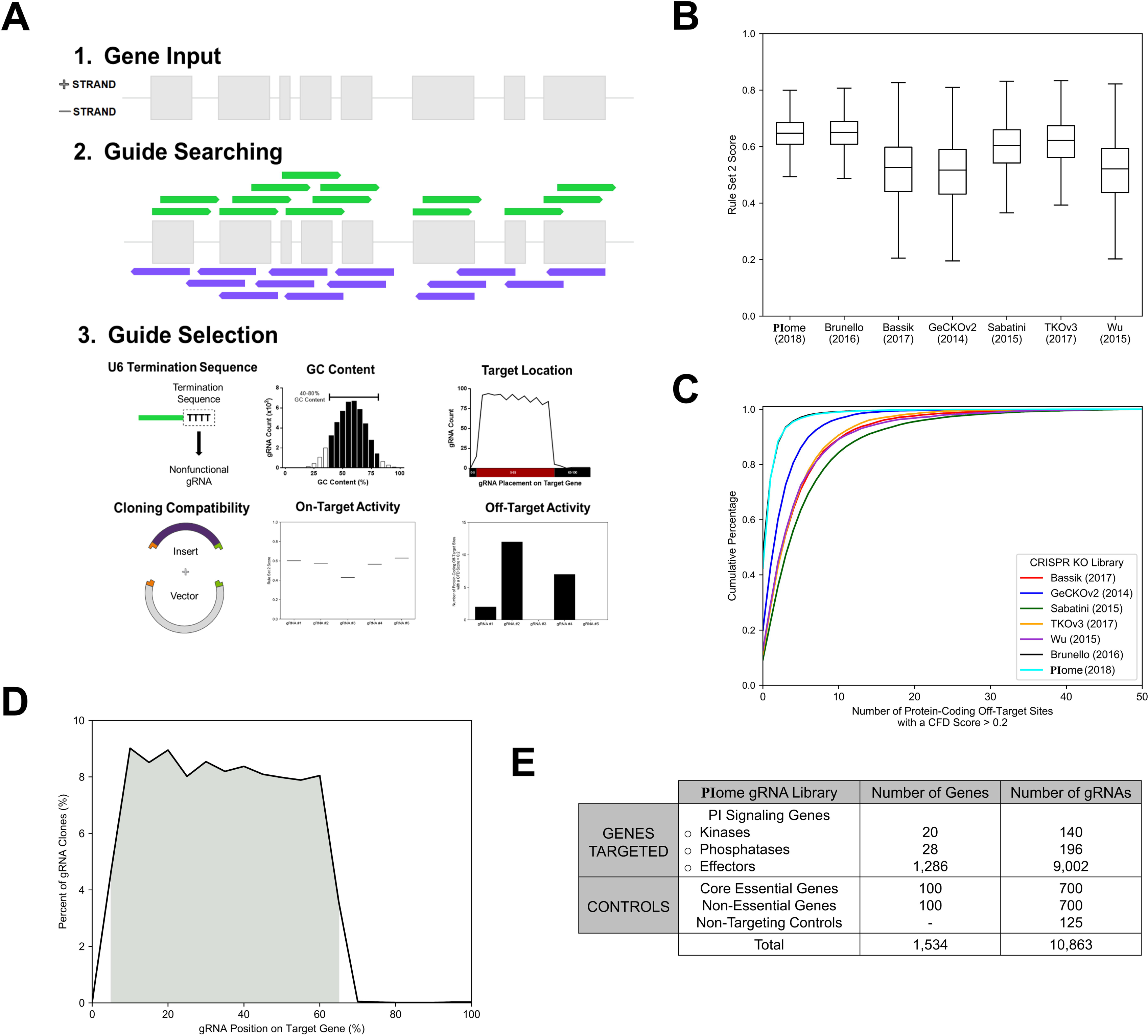
PI-focused CRISPR gRNA Library Design. (a) gRNAs for a given gene in the **PI**ome CRISPR KO library were designed using a three-step process: (1) Gene Input, (2) Guide Searching, and (3) Guide Selection (e.g. GC Content, Target Location). (b) Distribution of Rule Set 2 scores across CRISPR KO libraries. (c) Cumulative distribution plot of the number of protein-coding off-target sites with a CFD score > 0.2 across CRISPR KO libraries. (d) Percentage of gRNAs in the **PI**ome CRISPR KO library that target a given position on the CDS, as a percentage of gene length. The 5-65% CDS region is highlighted in grey. (e) Table of **PI**ome CRISPR KO library details.

Among the selection criteria, gRNA activity, specificity, and location were of utmost importance. We incorporated the latest Rule Set 2 and Cutting Frequency Determination (CFD) scoring metrics to select gRNAs that maximize on-target activity and minimize off-target effects.^19^ The **PI**ome library, alongside the Brunello library - a genome-wide library built on the same algorithm - is enriched for gRNAs predicted to be most active and specific compared to other CRISPR libraries (**Figure 1B-C**). Additionally, we preferred gRNAs targeting the 5’ end of the coding sequence (CDS) to increase the likelihood of generating a knockout (**Figure 1D**). The **PI**ome CRISPR KO library contains 10,738 unique gRNAs targeting 1,534 protein-coding genes, at 7 gRNAs per gene, and 125 non-targeting control gRNA (**Figure 1E**).

### PI-focused CRISPR-Cas9 Screen Identifies Novel Essential Genes in PANC-1

To identify essential PI-associated genes in PDAC, we performed a PI-focused CRISPR-Cas9 dropout screen in PANC-1, a human PDAC cell line. PANC-1 cells stably expressing Cas9 (**Figures S1A-B**) were transduced with the 11k **PI**ome CRISPR KO library at an MOI of 0.3 and 1000-fold coverage. After selection, cells were sampled at the initial timepoint (T0) and every 3 days, from day 3 (T03) until day 21 (T21), or ∼10 doublings. Genomic DNA was isolated at each timepoint, and gRNA abundance was measured by NGS (**Figure 2A** and **Table S2**).

**Figure 2.**
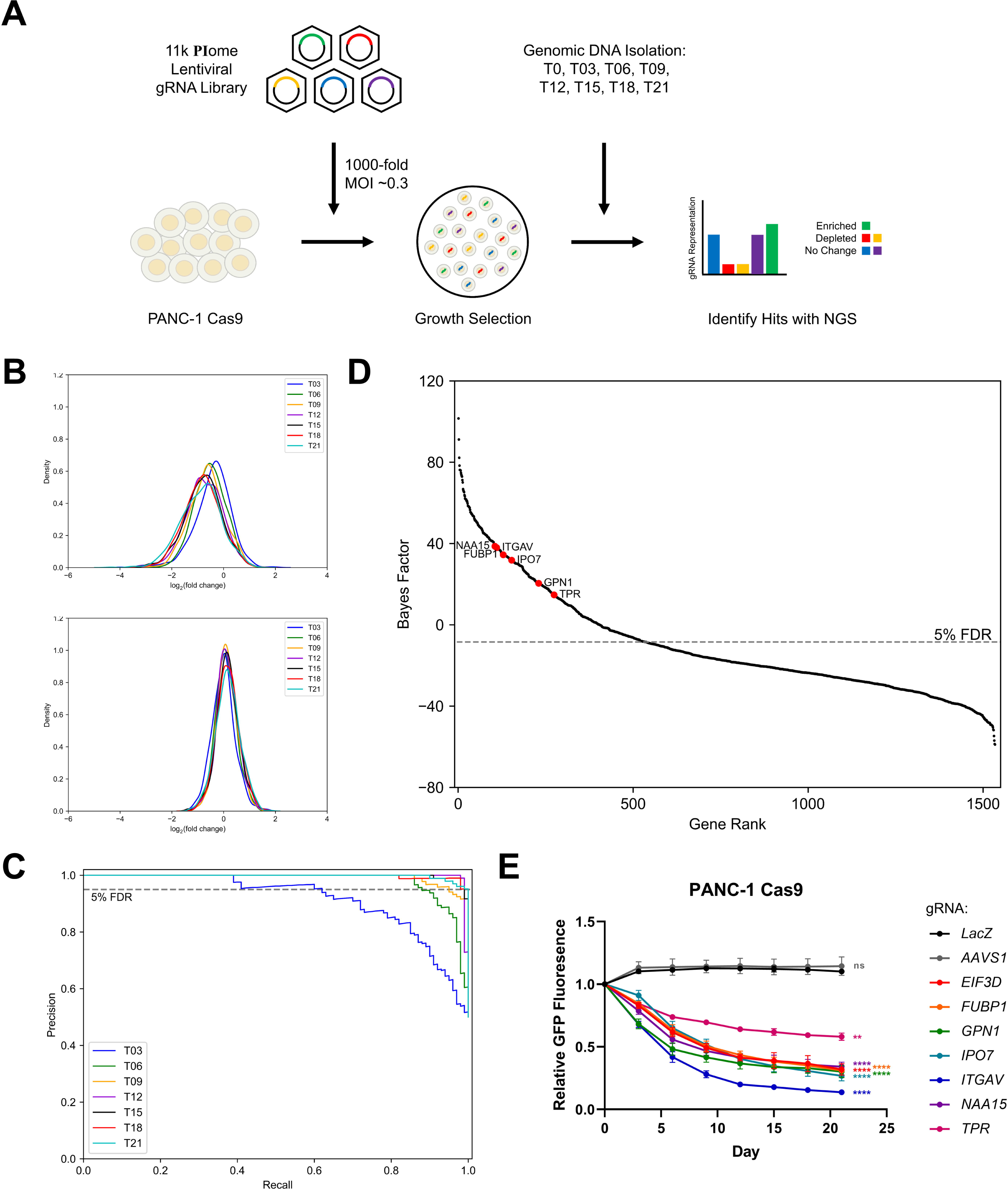
PI-focused CRISPR-Cas9 Screen Identifies Novel Essential Genes in PANC-1. (a) Schematic illustrates the workflow of the CRISPR-Cas9 dropout screen in PANC-1 cells. The 11k **PI**ome lentiviral gRNA library was transduced into Cas9-expressing PANC-1 cells at an MOI of 0.3 and 1000-fold coverage. Successfully transduced cells were selected with puromycin, split into three replicates, and passaged every three days at 1000-fold coverage. Cells were collected at the initial timepoint (T0) and every 3 days, from day 3 (T03) until day 21 (T21), for genomic DNA extraction. gRNA representation was determined by NGS and analyzed using the BAGEL algorithm. (b) Fold-change distributions of gRNA targeting essential genes (top) and nonessential genes (bottom) at the indicated time points in PANC-1. (c) Precision-recall curves from the PI-focused CRISPR-Cas9 screen in PANC-1. Values were generated using gold-standard sets of essential and nonessential genes ^11,12^. The dashed line denotes the 5% FDR threshold. (d) Ranked gene-level BF scores from the PI-focused CRISPR-Cas9 screen in PANC-1 at T21. Selected genes are indicated. The dashed line denotes the 5% FDR threshold. (e) Competition assay in PANC-1 Cas9 cells showing growth defects upon expression of gRNA targeting candidate hits. Each point represents the mean±SEM from n=3 independent experiments. ns p > 0.05, ** p < 0.01, **** p < 0.0001 from gRNA-*LacZ* by One-Way ANOVA with Dunnett’s Post-Hoc.

The fold-change distribution of gRNAs targeting essential genes was significantly left-shifted to those targeting nonessential genes, with this shift increasing over time, indicating that the screen functioned as designed (**Figure 2B**). We then used the BAGEL algorithm to calculate a Bayes Factor (BF) for each gene by comparing the gRNA fold-change to gold-standard sets of essential and nonessential genes (a high BF indicates increased confidence that the gene knockout results in a growth defect).^11,12,15^ Screen performance was further evaluated by calculating precision-recall curves at each timepoint, with later timepoints showing improved recall over earlier timepoints (**Figure 2C**).

A total of 106 genes were essential across all timepoints, at a minimum BF of 3 and 5% FDR, including 78 previously identified essential genes and 28 candidate essential genes (**Table 1**).^12^ Among the 28 candidate essential genes, only 6 genes (**Figure 2D**) exhibited statistically significant differences in survival when assessed in the TCGA-PAAD dataset. These include *FUBP1*, *GPN1*, *IPO7*, *ITGAV*, *NAA15*, and *TPR* (**Figures S1C-H**).

**Table 1.** Candidate Hits from PI-focused CRISPR Dropout Screen in PANC-1.

| PANC-1 |  |  |  |
| --- | --- | --- | --- |
| *TLN1* | *IPO7* | *ITGAV* | *TPR* |
| *ATP6V1B2* | *CWC22* | *RANBP1* | *NAA15* |
| *COPS5* | *STIP1* | *RNMT* | *SUPT16H* |
| *SRSF11* | *ACTR1A* | *DNM1L* | *ADRM1* |
| *MYH9* | *GPN1* | *DCTN2* | *RPL38* |
| *COPS8* | *LEMD2* | *RSL1D1* | *ZC3H18* |
| *SRP54* | *FUBP1* | *WNK1* | *FAM50A* |

To validate these hits, individual gene knockouts (two gRNAs per gene) were generated and tested in competitive cell growth assays (**Figure S1I**). Over a period of 21 days, PANC-1 Cas9 cells harboring gRNAs targeting the genes of interest were outcompeted by cells expressing the non-targeting control, gRNA-*LacZ* (**Figure 2E**). Taken together, these results indicate that our PI-focused CRISPR-Cas9 screen successfully identified essential genes whose expression levels are associated with a poor prognosis in patients with PDAC.

### PI-focused CRISPR-Cas9 Screen Identifies Novel Essential Genes in OCI-AML2

To identify novel essential genes in AML, we performed a PI-focused CRISPR- Cas9 dropout screen in OCI-AML2, a human AML cell line. OCI-AML2 cells stably expressing Cas9 (**Figure S2A-B**) were transduced with the 11k **PI**ome CRISPR KO library at an MOI of 0.3 and 1000-fold coverage. After selection, cells were sampled at the initial timepoint (T0), day 7, and every 3 days, from day 10 (T03) until day 22 (T15), or ∼12 doublings. Genomic DNA was isolated at each timepoint, and gRNA abundance was measured by NGS (**Figure 3A** and **Table S3**).

**Figure 3.**
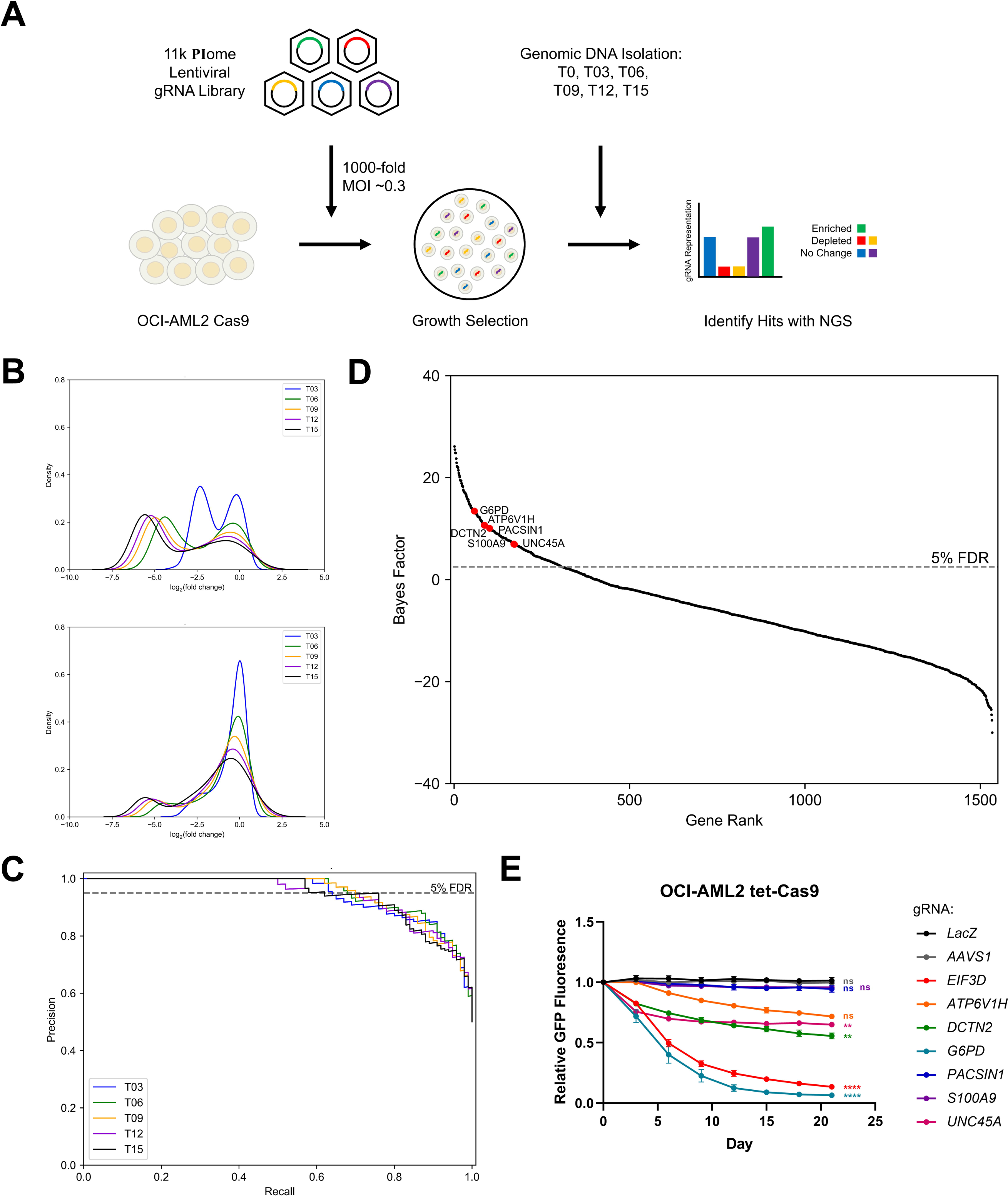
PI-focused CRISPR-Cas9 Screen Identifies Novel Essential Genes in OCI- AML2. (a) Schematic illustrates the workflow of the CRISPR-Cas9 dropout screen in OCI- AML2 cells. The 11k **PI**ome lentiviral gRNA library was transduced into Cas9-expressing OCI-AML2 cells at an MOI of 0.3 and 1000-fold coverage. Successfully transduced cells were selected with puromycin, split into three replicates, and passaged every three days at 1000-fold coverage. Cells were collected at the initial timepoint (T0), day 7, and every 3 days, from day 10 (T03) until day 22 (T15), for genomic DNA extraction. gRNA representation was determined by NGS and analyzed using the BAGEL algorithm. (b) Fold-change distributions of gRNA targeting essential genes (top) and nonessential genes (bottom) at the indicated timepoints in OCI-AML2. (c) Precision-recall curves from the PI-focused CRISPR-Cas9 screen in OCI-AML2. Values were generated using gold-standard sets of essential and nonessential genes ^11,12^. The dashed line denotes the 5% FDR threshold. (d) Ranked gene-level BF scores from the PI-focused CRISPR-Cas9 screen in OCI-AML2 at T15. Selected genes are indicated. The dashed line denotes the 5% FDR threshold. (e) Competition assay in OCI-AML2 tet-Cas9 cells showing growth defects upon expression of gRNA targeting select candidate hits. Each point represents the mean±SEM from n=3 independent experiments. ns p > 0.05, ** p < 0.01, **** p < 0.0001 from gRNA-*LacZ* by One-Way ANOVA with Dunnett’s Post-Hoc.

The fold-change distribution of gRNAs targeting essential genes was significantly left-shifted to those targeting nonessential genes, with this shift increasing over time, indicating that the screen functioned as designed (**Figure 3B**). We then used the BAGEL algorithm to calculate a BF for each gene by comparing the gRNA fold-change to gold standard sets of essential and nonessential genes.^11,12,15^ Using the gold-standard reference sets, screen performance was further evaluated by calculating precision-recall curves at each timepoint, with all timepoints showing similar recall (**Figure 3C**).

A total of 174 essential genes were essential across all timepoints, at a minimum BF of 3 and 5% FDR, including 90 previously identified essential genes and 84 candidate essential genes (**Table 2**).^12^ Surprisingly, out of the 84 candidate essential genes, only 6 genes (**Figure 3D**) exhibited statistically significant differences in survival in the TCGA-LAML dataset: *ATP6V1H*, *DCTN2*, *G6PD*, *PACSIN1*, *S100A9,* and *UNC45A* (**Figure S2C-H**).

**Table 2.** Candidate Hits from PI-focused CRISPR Dropout Screen in OCI-AML2.

| OCI-AML2 |  |  |  |
| --- | --- | --- | --- |
| *CD44* | *ARF4* | *PCBP1* | *UHRF1* |
| *RPTOR* | *TOMM34* | *PLEKHA3* | *LIMS1* |
| *CACNA1C* | *ARHGEF4* | *ATP2B2* | *WNK1* |
| *SLC25A3* | *NPM1* | *PMVK* | *TPR* |
| *DDX5* | *G6PD* | *SNX13* | *LRCH3* |
| *CBFB* | *PPP2R2A* | *PTPMT1* | *NFIC* |
| *COX6C* | *GADD45GIP1* | *PLEKHA4* | *SERBP1* |
| *WDR11* | *PRKD1* | *XPO4* | *WASHC2C* |
| *RPL7A* | *SRP54* | *TALDO1* | *SUPT16H* |
| *CAND1* | *VSNL1* | *RBM39* | *OR1S2* |
| *EPN3* | *TRPC6* | *STX7* | *MYO1D* |
| *PTPN11* | *NDUFB4* | *CCDC22* | *MTMR3* |
| *CLDN3* | *OSBP* | *PTER* | *PTPN9* |
| *PACSIN1* | *ZW10* | *PIP5K1B* | *UNC45A* |
| *PPIL4* | *MTMR2* | *PRKDC* | *RPL38* |
| *CNGA3* | *MCF2* | *TAS2R1* | *SMG1* |
| *TNPO1* | *SYNGR2* | *NACA* | *ARHGAP23* |
| *ATP6V1H* | *DOK4* | *IPMK* | *EEA1* |
| *PLCD1* | *EIF4E2* | *EFR3A* | *RPS6KC1* |
| *COPG1* | *PHETA2* | *CD46* | *CHMP4B* |
| *WDFY2* | *KLC1* | *DCTN2* | *S100A9* |

To validate candidate hits, individual gene knockouts (two gRNAs per gene) were generated and tested in competitive cell growth assays. Over a period of 21 days, OCI- AML2 tet-Cas9 cells harboring gRNAs targeting *DCTN2*, *G6PD*, and *UNC45A* were outcompeted by cells expressing the non-targeting control, gRNA-*LacZ* (**Figure 3E**). Taken together, these results indicate that our PI-focused CRISPR-Cas9 screen successfully identified essential genes whose expression levels are associated with a poor prognosis in patients with AML.

### PI-focused CRISPR-Cas9 Screen Identifies Hits Missed in Genome-wide CRISPR-Cas9 Screen

To evaluate sub-library performance, we compared the fitness scores of PI- focused CRISPR-Cas9 hits with those from genome-wide screens performed by other groups.^21^ In the PANC-1 screen, all but one (*FUBP1*) of the validated hits were found to be essential in both the PI-focused and genome-wide screen (**Figure 4A**). In the OCI- AML2 screen, two of the validated hits, *G6PD* and *UNC45A*, were not identified as essential in the genome-wide screen, while *ATP6V1H* trended towards significance in the competition assay (**Figure 4B**). Notably, *PACSIN1* and *S100A9* were validated as non-essential, exhibiting opposite results between the PI-focused and genome-wide screen (**Figure 4B**). Among all the hits, 4 of the 28 were not identified as essential in the PANC- 1 genome-wide screen, while 51 of the 80 (with data missing for four) were not identified as essential in the OCI-AML2 genome-wide screen. All in all, these results underscore the value of sub-libraries in uncovering novel genetic vulnerabilities that may be missed by genome-wide approaches.

**Figure 4.**
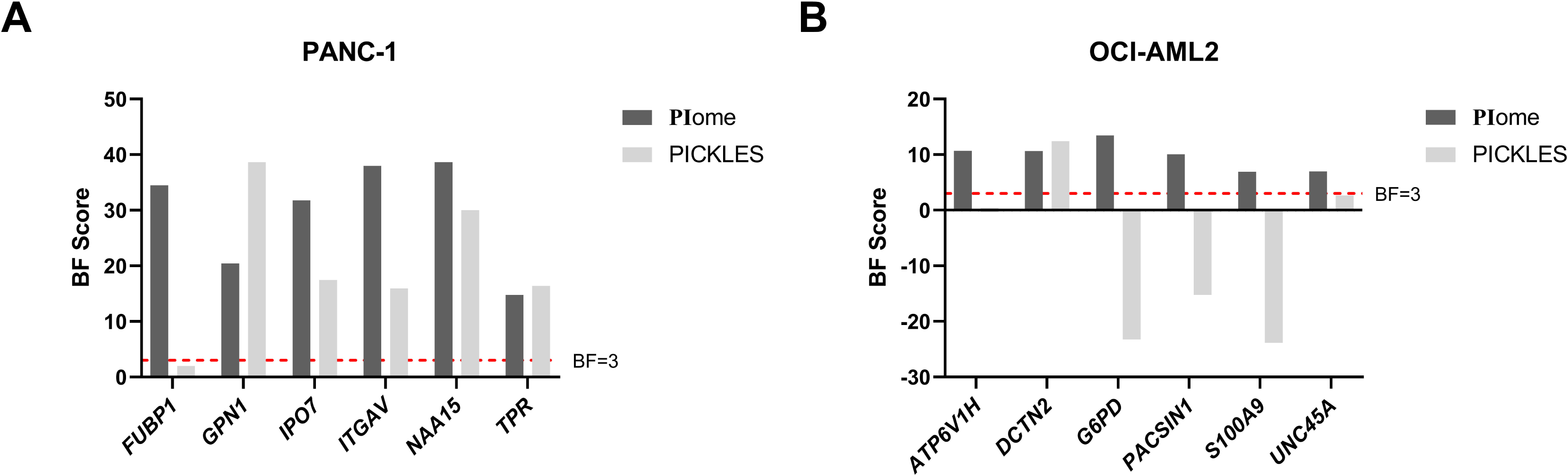
PI-focused CRISPR-Cas9 Screen Identifies Hits Missed in Genome-wide CRISPR-Cas9 Screen. Comparison of candidate hit fitness scores between the PI- focused CRISPR screen and a genome-wide CRISPR screen from the PICKLES database ^21^ in (a) PANC-1 and (b) OCI-AML2 cells. The dashed line denotes the BF=3 cutoff.

### PI-focused CRISPR Screen Identifies Novel Regulators of Colony Formation in OCI- AML2

To identify regulators of colony formation, a surrogate measure of leukemic stem cell (LSC) stemness *in vitro*, we performed a PI-focused CRISPR-Cas9 colony-forming unit (CFU) screen in OCI-AML2 cells. OCI-AML2 cells stably expressing Cas9 were transduced with the 11k **PI**ome CRISPR KO library at an MOI of 0.3 and 500-fold coverage. After selection, cells were sampled at the initial timepoint (T0), day 7, and plated in colony-forming cell assays for 7 days. Cells with the desired phenotype were enriched by solubilizing and repeating the procedure two times. Genomic DNA was isolated at each timepoint, and gRNA abundance was measured by NGS (**Figure 5A** and **Table S4**).

**Figure 5.**
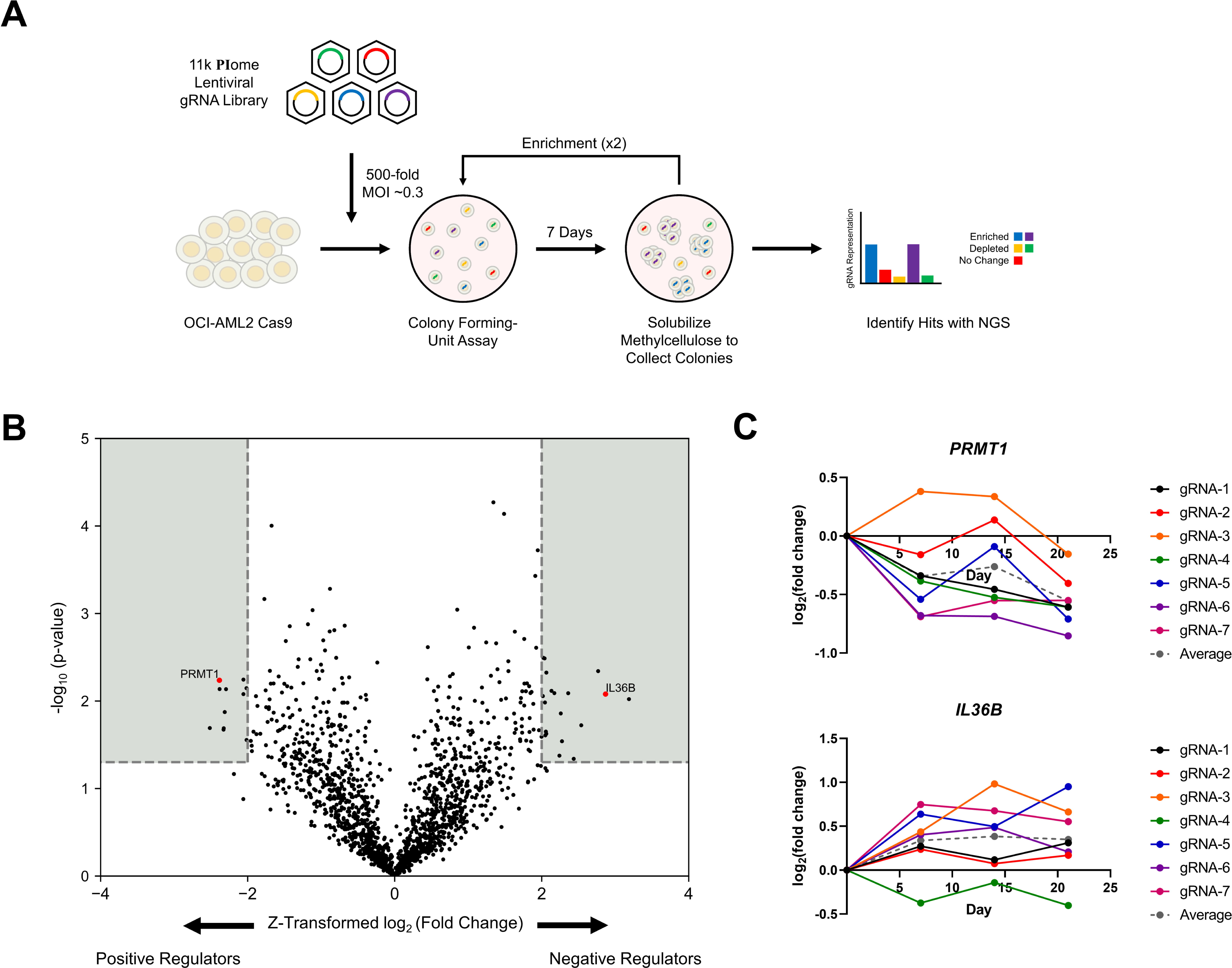
PI-focused CRISPR-Cas9 Screen Identifies Novel Regulators of Colony Formation in OCI-AML2. (a) Schematic illustrates the workflow of the CRISPR-Cas9 CFU screen in OCI-AML2 cells. The 11k **PI**ome lentiviral gRNA library was transduced into Cas9-expressing OCI-AML2 cells at an MOI of 0.3 and 500-fold coverage. Successfully transduced cells were selected with puromycin, sampled at the initial timepoint (T0, day 7), and plated in methylcellulose colony-forming cell assays for 7 days. Cells with the desired phenotype were enriched by solubilizing the methylcellulose and repeating the procedure two times. Genomic DNA was extracted at each timepoint, and gRNA abundance was determined by NGS. (b) Volcano plot displaying fold change versus significance for each gene tested in the PI-focused CRISPR gRNA library. A quadrant (light gray) bounded by |Z-transformed log_2_ (fold change)| > 2 and -log_10_ (p-value) < 0.05 contains select significant hits. (c) Fold change of gRNAs targeting *PRTM1* (top) and *IL36B* (bottom) in the PI-focused CRISPR-Cas9 CFU screen.

A total of 28 regulators of colony formation were identified, comprising 17 negative regulators and 11 positive regulators, based on the following criteria: 1) |Z-transformed log_2_ (fold change)| > 2, 2) p-value < 0.05, and 3) at least 4 functional gRNAs (**Figure 5B** and **Table 3**). Notably, there was no overlap between the essential genes identified and the negative regulators of colony formation, suggesting that the identified genes may play distinct roles in LSC maintenance and differentiation. For instance, PRMT1, a primary type I arginine methyltransferase, was previously identified in various studies as a positive regulator of colony formation, demonstrating reduced colony growth upon *PRMT1* knockdown (**Figure 5B-C**).^22–24^ In contrast, IL36B, a cytokine of the interleukin 1 family, was shown to be enriched in LSC-depleted cells, suggesting its role as a negative regulator of colony formation (**Figure 5B-C**).^25^ Together, these findings from the novel CFU screen identify crucial regulators of colony formation, providing important insights into the mechanisms governing LSC maintenance and differentiation.

**Table 3.**
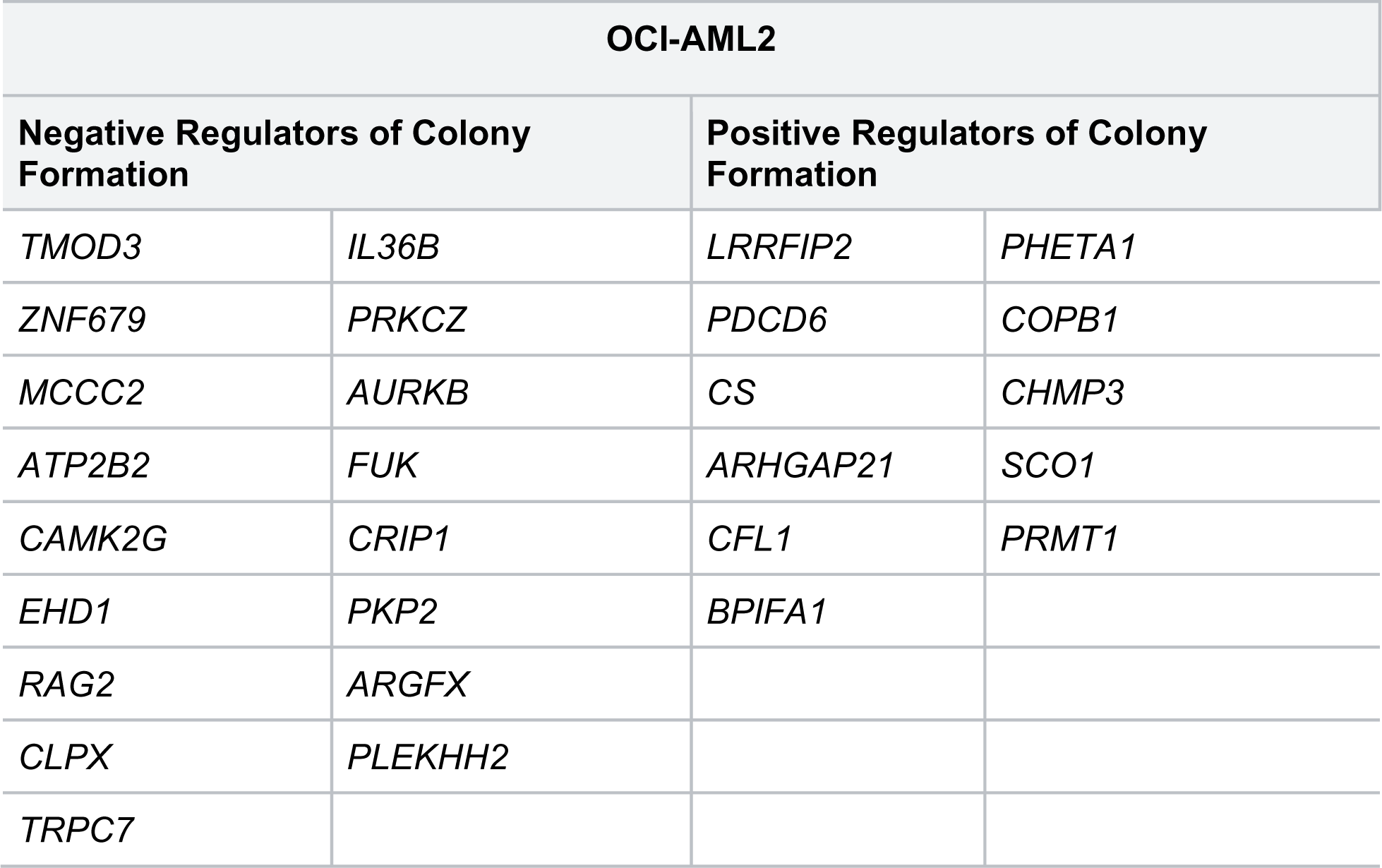
Candidate Hits from PI-focused CFU Screen in OCI-AML2.

| OCI-AML2 |  |  |  |
| --- | --- | --- | --- |
| Negative Regulators of Colony Formation |  | Positive Regulators of Colony Formation |  |
| *TMOD3* | *IL36B* | *LRRFIP2* | *PHETA1* |
| *ZNF679* | *PRKCZ* | *PDCD6* | *COPB1* |
| *MCCC2* | *AURKB* | *CS* | *CHMP3* |
| *ATP2B2* | *FUK* | *ARHGAP21* | *SCO1* |
| *CAMK2G* | *CRIP1* | *CFL1* | *PRMT1* |
| *EHD1* | *PKP2* | *BPIFA1* |  |
| *RAG2* | *ARGFX* |  |  |
| *CLPX* | *PLEKHH2* |  |  |
| *TRPC7* |  |  |  |

## Discussion

PI homeostasis and functions are governed by an intricate interplay between lipids, kinases, phosphatases, and effectors. This coordination is crucial for the regulation of many essential cellular processes. Previous work by other groups has focused on the role of PI-metabolizing enzymes in cell growth.^8,9^ However, to our knowledge, there has not been a comprehensive investigation into the complete PI signaling system, which includes a broad examination of kinases, phosphatases, and effectors involved in the process. In this study, we developed and employed a novel PI-focused CRISPR gRNA library to conduct genetic screens in PDAC and AML.

Using our PI-focused CRISPR gRNA library, we performed dropout screens in PANC-1 and OCI-AML2, models of PDAC and AML, respectively. We identified 28 essential genes in PANC-1 and 84 essential genes in OCI-AML2. Among the hits, there were 5 genes that overlapped between the two screens. This disparity highlights the unique molecular mechanisms and pathways that underlie the development of each cancer type. Among the hits identified, we specifically chose genes with expression levels linked to poor prognosis for further individual validation. This selection process highlights their utility as prognostic biomarkers and potential targets for therapeutic interventions. From the PANC-1 screen, only *IPO7* and *ITGAV* have been shown to play a role in PDAC pathogenesis, while the other identified genes (*FUBP1*, *GPN1*, *NAA15*, *TPR*) have been implicated in the pathogenesis of different cancers ^26–44^. Similarly, in the OCI-AML2 screen, only *G6PD* has been implicated in AML pathogenesis, while the other identified genes (*DCTN2*, *UNC45A*) have been associated with different types of cancer.^45–58^ Together, these findings highlight context-dependent genetic vulnerabilities and identify potential prognostic biomarkers for future investigation in PDAC and AML.

Given the large number of publicly available CRISPR-Cas9 screening data, we compared the fitness scores of candidate hits from our PI-focused CRISPR-Cas9 screens with those from genome-wide screens conducted by other groups. This comparison allowed us to identify potential hits that were overlooked in the broader screens, thereby validating the specificity and effectiveness of our PI-focused approach. Using the PICKLES database, we identified and validated *FUBP1*, *G6PD*, *UNC45A*, and possibly *ATP6V1H* as essential genes that would likely be missed in genome-wide screens.^21^ Looking at all candidate hits, only 4 of 28 essential genes identified in the PANC-1 screen were missed in genome-wide screens. In contrast, a notable 51 of 80 (with data missing for four) essential genes identified in the OCI-AML2 screen were missed in the genome-wide screens, underscoring the added value of focused libraries in identifying genetic vulnerabilities.

The advantages of using focused libraries, such as the PI-focused library we developed, include increased coverage and resolution.^10,59–62^ By increasing the number of gRNAs per gene, we achieved more comprehensive coverage, reducing the likelihood of false negatives that occur when genes are poorly represented in genome-wide screens. Additionally, enhanced resolution, exemplified by the increased number of cells expressing a given gRNA, enables the identification of subtle yet significant effects of gene function that may not be captured in genome-wide screens. However, it’s important to note that despite the advantages of focused libraries, our approach also has its limitations. For instance, the identification of potential hits, such as *PACSIN1* and *S100A9*, in our screen that were not validated in the competition assay highlights the possibility of false positives or subtle effects that may have not been detected under competitive conditions. Thus, our PI-focused CRISPR-Cas9 screening approach offers valuable insights into the genetic vulnerabilities of PDAC and AML, particularly in uncovering potential false negatives and subtle effects that may be missed in genome-wide screens.

Another advantage of focused libraries is the ability to perform screens across a wider range of screening conditions. One commonly used positive-selection screen is the transwell migration or invasion assay, which is aimed at identifying genes implicated in either cell migration or invasion.^63–67^ While this is possible with genome-wide libraries, certain screening conditions may not be feasible due to the vast number of targets and the complexity involved. One example is the CFU assay, which assesses the ability of single cells to form a colony, serving as a surrogate measure of stemness in leukemia. Some important considerations in the assay include a low cell number to ensure colonies originate from single cells, avoiding stacking more than two plates in the incubator to prevent vibrations from destroying the colonies, and carefully preparing the FBS- methylcellulose-cell mixture to ensure proper distribution when mixing. Given these constraints, genome-wide screens may be unfeasible under all these conditions, making a focused library a better option.

In our PI-focused CFU screen, we identified 28 regulators of colony-formation, comprising 17 negative regulators and 11 positive regulators, using three main criteria: fold-change cutoff, statistical significance, and at least four functional gRNAs. Interestingly, there was no overlap between the essential genes identified in the negative-selection screen and the negative regulators identified in the positive-selection screen, highlighting the utility of our approach for identifying genes with distinct roles in LSC maintenance and differentiation. Among the hits identified, *IL36B* and *PRMT1* stood out as genes with known roles in leukemic colony formation.^22–25^ However, little is known about the remaining genes, with some (*CAMK2G*, *CRIP1*, *EHD1*) being linked to stemness in other cancers..^68–74^ Therefore, our innovative CFU screening method effectively pinpointed genes already recognized for their involvement in stemness, thereby bolstering the credibility of the assay. This outcome also paves the way for additional explorations into previously unexplored genes and their potential implications in leukemia.

In summary, by utilizing a PI-focused CRISPR gRNA library, we discovered previously unidentified genes as being essential in PDAC and AML. We show that focused libraries have the power to uncover false negatives, detect subtle effects, and adapt to different screening conditions. This efficient and targeted approach allows for more comprehensive and nuanced investigations into the genetic underpinnings of cancer, paving the way for the discovery of new therapeutic targets and biomarkers.

## Acknowledgements

Dr. Leonardo Salmena is the recipient of a Tier II Canada Research Chair (CRC) and supported through the Human Frontier Career Development Program (HFSP) award. Funding for this research was provided in part by the Temerty Faculty of Medicine and Department of Pharmacology and Toxicology, University of Toronto and awards from the Canada Foundation for Innovation (CFI-33505) and the Canadian Institutes for Health Research (Project Grant #399716, #480743).

## Author Contributions

Daniel KC Lee - Conceptualization, Methodology, Software, Validation, Formal Analysis, Investigation, Data Curation, Visualization, Project Administration, Writing - original draft, Writing - review & editing

Ryan Loke - Formal Analysis, Investigation

Jonathan TS Chow - Formal Analysis, Investigation

Martino M Gabra - Software

Leonardo Salmena - Conceptualization, Investigation, Resources, Supervision, Project Administration, Funding Acquisition, Writing - original draft, Writing - review & editing

## Abbreviations

PIs: Phosphoinositides
PDAC: Pancreatic Ductal Adenocarcinoma
AML: Acute Myeloid Leukemia
CRISPR: Clustered Regularly Interspaced Short Palindromic Repeats
CFD: Cutting Frequency Determination
CDS: Coding Sequence
BAGEL: Bayesian Analysis of Gene Essentiality

---

**Figure S1. - Related to Figure 2. PANC-1 CRISPR-Cas9 Dropout Screen Supporting Information.** (a) Western blot of Cas9 levels in parental and clonal Cas9-expressing PANC-1 cells. (b) Flow cytometry profiles of EGFP levels in parental and clonal Cas9- expressing PANC-1 cells following expression of EGFP and a gRNA targeting EGFP. Kaplan-Meier plots for (c) *FUBP1*^high^ vs. *FUBP1*^low^, (d) *GPN1*^high^ vs. *GPN1*^low^, (e) *IPO7*^high^ vs. *IPO7*^low^, (f) *ITGAV*^high^ vs. *ITGAV*^low^, (g) *NAA15*^high^ vs. *NAA15*^low^, and (h) *TPR*^high^ vs. *TPR*^low^ PDAC patient overall survival in the TCGA-PAAD dataset. (i) Schematic illustrates the competitive cell growth assay used to validate candidate essential genes. Cas9 cells are first infected and selected for a gRNA targeting *LacZ* (GFP-) or a gRNA targeting candidate essential genes (GFP+). Cells are then mixed 1:1 and cultured over a period of 21 days with GFP monitoring every 3 days. gRNA targeting essential genes will drop out of the population (decrease in GFP%), whereas gRNA targeting nonessential genes will be maintained in the population (no change in GFP%).

**Figure S2. - Related to Figure 3. OCI-AML2 CRISPR-Cas9 Dropout Screen Supporting Information.** (a) Western blot of Cas9 levels in parental and clonal Cas9- expressing OCI-AML2 cells. (b) Flow cytometry profiles of EGFP levels in parental and clonal Cas9-expressing OCI-AML2 cells following expression of EGFP and a gRNA targeting EGFP. Kaplan-Meier plots for (c) *ATP6V1H*^high^ vs. *ATP6V1H*^low^, (d) *DCTN2*^high^ vs. *DCTN2*^low^, (e) *G6PD*^high^ vs. *G6PD*^low^, (f) *PACSIN1*^high^ vs. *PACSIN1*^low^, (g) *S100A9*^high^ vs. *S100A9*^low^, and (h) *UNC45A*^high^ vs. *UNC45A*^low^ AML patient overall survival in the TCGA-LAML dataset.

