## Supplementary figures and images for "Phosphoinositide-focused CRISPR screen identifies novel genetic vulnerabilities in PDAC and AML cells"

### Supplemental Figure 1

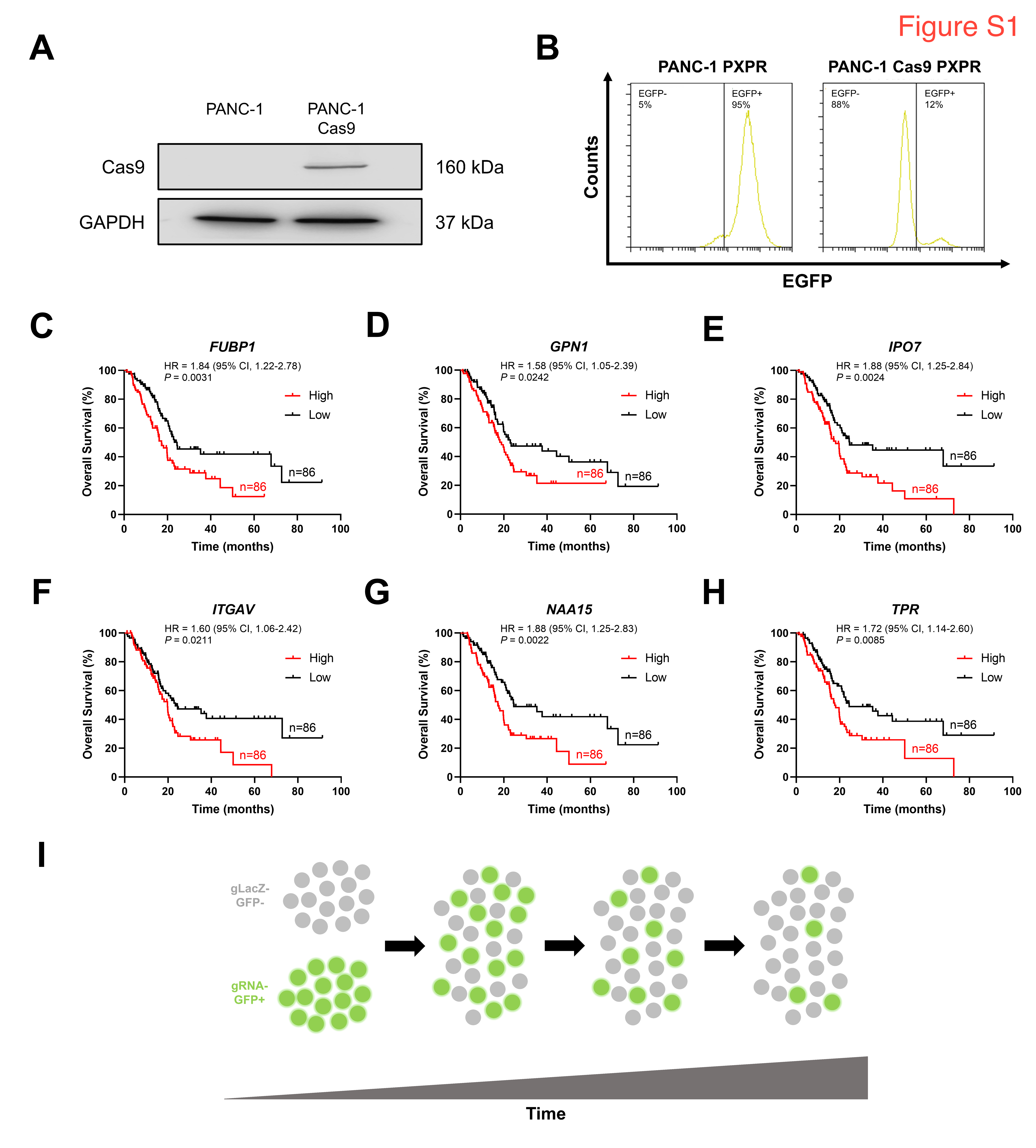

### Supplemental Figure 2

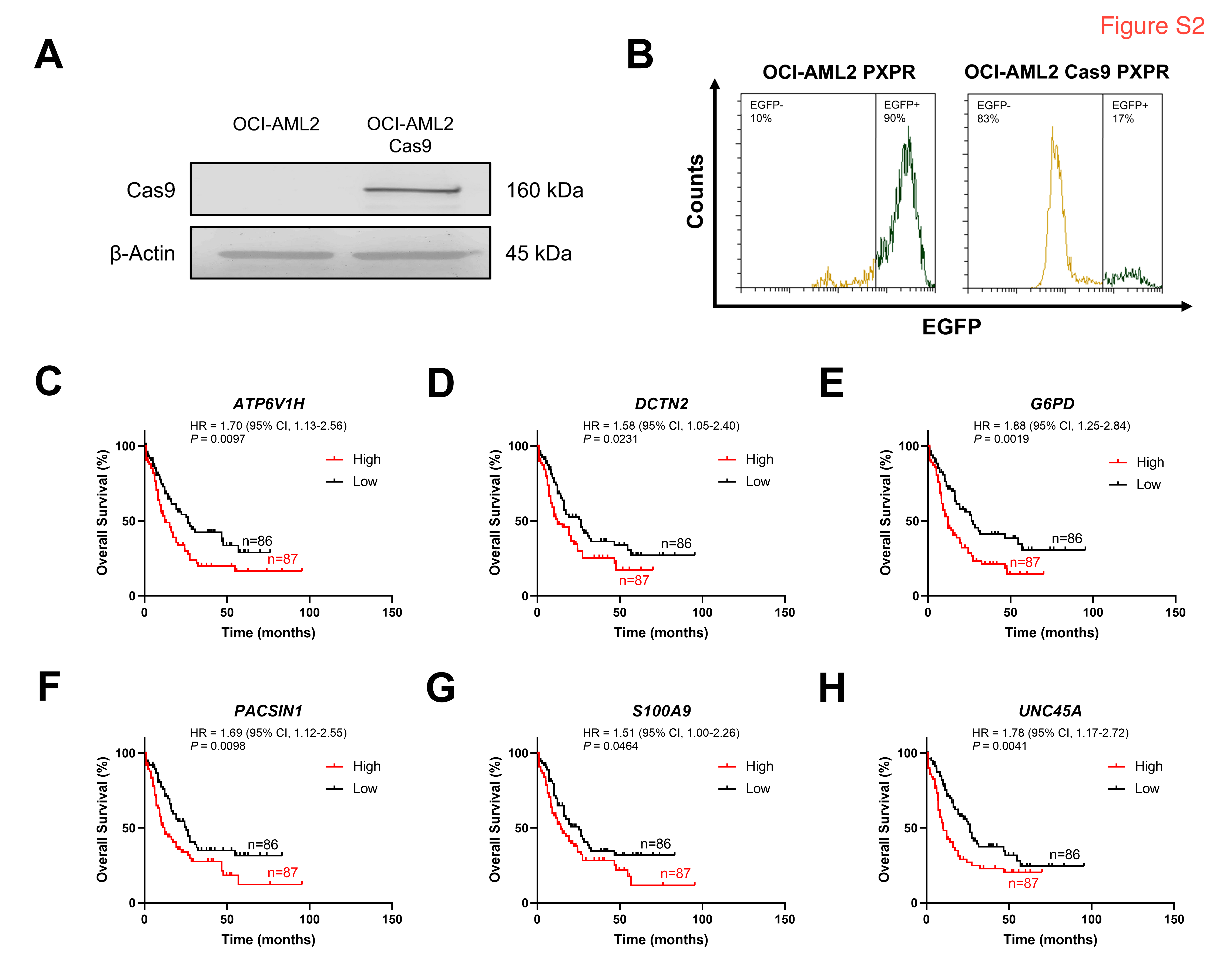
